# A two-stage diffusion modeling approach to the compelled-response task

**DOI:** 10.1101/2020.05.22.110288

**Authors:** Adele Diederich, Hans Colonius

**Affiliations:** Health: Life Sciences & Chemistry, Jacobs University, Bremen, Germany; Department of Psychology and Research Center Neurosensory Science, Carl von Ossietzky Universität, Oldenburg, Germany

**Keywords:** compelled-response task, race-to-threshold model, two-stage diffusion model, non-time-homogeneity

## Abstract

The issue of how perception and motor planning interact to generate a given choice between actions is a fundamental question in both psychology and neuroscience. Salinas and colleagues have developed a behavioral paradigm, the compelled-response task, where the signal that instructs the subject to make an eye movement is given before the cue that indicates which of two possible target choices is the correct one. When the cue is given rather late, the participant must guess and make an uninformed random choice. Perceptual performance can be tracked as a function of the amount of time during which sensory information is available. In Salina’s accelerated race-to-threshold model, two variables race against each other to a threshold, at which a saccade is initiated. The source of random variability is in the initial state of information buildup across trials. This implies that incorrect decisions are due to the inertia of the racing variables that have, at the start, sampled a constant buildup in the “wrong” direction. Here we suggest an alternative, non-time-homogeneous two-stage-diffusion model that is able to predict both response time distributions and choice probabilities with a few easy-to-interpret parameters and without assuming cross-trial parameter variability. It is falsifiable at the level of qualitative features already, e.g. predicting bimodal RT distributions for particular gap times. It connects the compelled-response paradigm with an approach to decision making that has been uniquely successful in describing both behavioral and neural data in a variety of experimental settings for the last 40 years.

---

Riding your bike in a busy city street, you constantly have to predict events and prepare for quick decisions. Will the pedestrian entering the road actually stop to let you drive past? Will the door of the car you are just about to pass open suddenly? In both cases, you must decide either to go straight on or swerve. Your response will be the result of a perceptual decision, but how long did it take you prepare the movement and to make the decision before executing your motor response? Disentangling these different components of the reaction time (RT) has proven to be a challenging task even under highly controlled laboratory conditions. Within a psychophysical paradigm only the total amount of RT is directly observable, but even when underlying neural circuits are accessible, the task is not straightforward because neurons that encode perceptual decisions are often also involved in motor planning (e.g., Horwitz & Newsome, 1999). Over the last 150 years, experimental psychology has developed a host of paradigms to unravel the elemental processes involved in generating a response in such tasks within a certain amount of time (see Luce, 1986, for an overview).

More recently Salinas and colleagues have developed a paradigm that seems particularly suited in separating the process of perceptual decision making from motor planning and execution (Stanford, Shankar, Massoglia, Costello, & Salinas, 2010; Salinas, Shankar, Costello, Zhu, & Stanford, 2010; Salinas, Scerra, Hauser, Costello, & Stanford, 2014; Shankar et al., 2011). In this <u>compelled-saccade</u> (CS) <u>task</u> participants start fixating a spot displayed in the middle of the screen, and the color of the spot, red or green, indicates the target color. Then, two yellow spots –potential targets– appear equidistant to the left and right of the fixation point. Next, the fixation point disappears telling the participant to initiate a saccade (go signal). At this time, the position of target and distracter are not yet revealed. After a delay, the peripheral spots change color: one turns red and the other green. The delay between the offset of the fixation point (go screen) and the onset of the cue screen is called <u>gap time</u> and typically varies between 50 and 250 ms in steps of 25 ms. Thus, crucially, the signal that instructs the subject to start an eye movement, i.e. the disappearance of the fixation point, is given prior to the cue (color identity) and indicates which of two possible choices is the correct one, the target. When the cue is given rather late (or, say, never), the participant must guess and make an uninformed random choice. In contrast, the earlier the cue is given the more informed the participant will be in making a choice. The idea is that the motor process is initiated early on and that perceptual information, once presented, influences a motor plan that is already evolving. Probing the paradigm on monkeys, it has been suggested that perceptual performance can be tracked as a function of the amount of time during which sensory information is available (called <u>tachometric curve</u>) independently of motor demands (Stanford et al., 2010; Costello, Zhu, Salinas, & Stanford, 2013; Shankar et al., 2011). Fig 1 shows the timeline of one trial (cf. Stanford et al., 2010, Fig 1).

**Figure 1.**
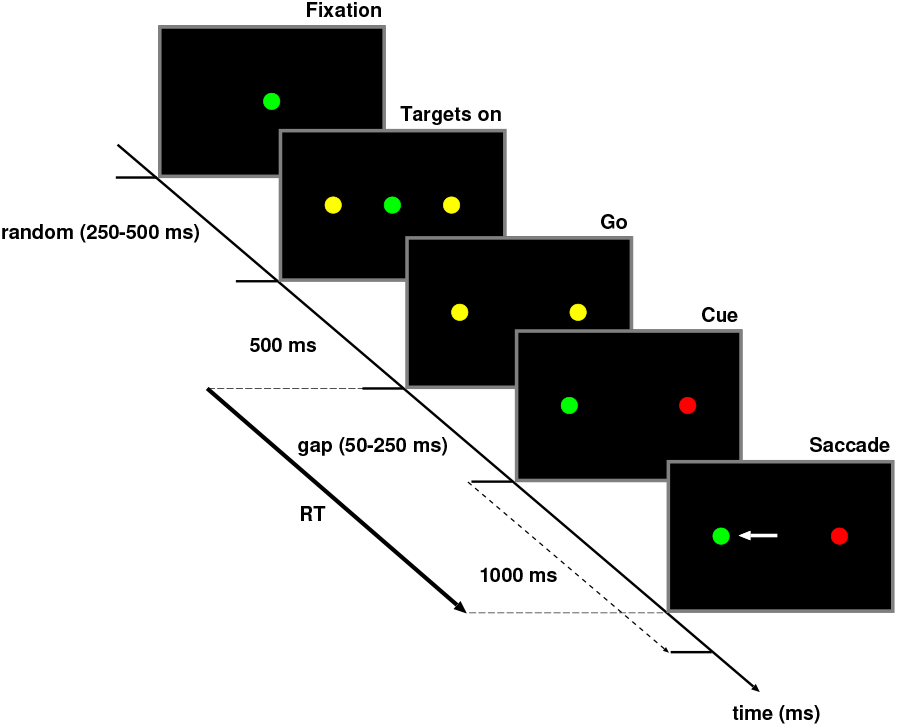
Compelled-saccade task. Timeline of events in the compelled-saccade task. The fixation point indicates the color of the target (green, in this example). The participant is instructed to gaze left or right as soon as the fixation point disappears (go screen). Target and distracter colors and positions are only revealed after a gap of 50–250 ms (onset of cue screen). A trial is correct if the participant makes an eye movement to the peripheral location that matches the color of the fixation point (green, in this example). Response time (RT) is defined from offset of the fixation point to saccade initiation.

Salinas and Stanford also proposed a computational model of perceptual decision making for the compelled-response task. Specifically, in their <u>accelerated race-to-threshold model</u> two variables race against each other to a threshold, at which a saccade is initiated. Each variable represents the motor plan to perform a saccade to one of the two spots, and the variable that first crosses the threshold determines the spot to which the saccade goes. Importantly, once the target position is signaled, the incoming perceptual information differentially modulates their trajectories toward it: the variable corresponding to the correct target accelerates and the one corresponding to the incorrect distractor decelerates. This model was fit successfully to data from two monkeys trained in this task. The authors showed that it captured both the full response time distributions and the dependence of decision accuracy on the effective sensory processing time. Specifically, they concluded that stimulus information (color identification) needs to be processed for just 25–50 ms to have an impact on the developing oculomotor plan.

Nevertheless, as observed by Drugowitsch and Pouget (Drugowitsch & Pouget, 2010) in a comment on the Stanford et al. (2010) paper, the accelerated race-to threshold model has an unusual feature: “*all stochasticity in the subject’s responses is attributed to the random choice of the initial race speeds.* […] *No extra noise is added at a later time, even after the delay period, when the color of the target is revealed. As a result, incorrect choices occur only because of the inertia of the racing variables.”* They also state that this is in contrast to standard models of decision making, such as diffusion or race models where response time variability is due to sensory noise and uncertainty in the stimulus itself, and suggest that *“it would be interesting to see whether the drift diffusion model and its neural counterparts would fit the data of these experiments as well as the deterministic race model…”* (see Drugowitsch & Pouget, 2010, p. 280).

Taking up the proposal by Drugowitsch and Pouget, here we propose an alternative to the race-to-threshold model, a <u>two-stage diffusion model</u> of behavior in the compelled response task. Note that we do not claim this alternative modeling approach –limited as it is here to behavioral data– to be uniquely preferable to the accelerated race-to threshold model. Rather, as a proof of principle, we show that stimulus processing in this task can be accounted or by an approach that (i) has been uniquely successful in describing the time-course of behavioral decision processes (RT and accuracy) (e.g., Ratcliff, Smith, Brown, & McKoon, 2016; Diederich, 1992; Ricciardi, 1977) and that (ii) has been shown to match the pattern of activity growth recorded in SC buildup neurons (Ratcliff, Cherian, & Segraves, 2003; Smith & Ratcliff, 2004). Importantly, we will point out in the final section what our demonstration contributes to the current discussion in the general area of race and diffusion modeling in psychology.

This note is organized as follows. After adding some details about the race-to-threshold model, we present the two-stage diffusion model in a non-technical manner and consider different scenarios for the compelled-response task. Various predictions of the two-stage diffusion model, hereafter abbreviated as 2SD model, are explicated in the two subsequent sections. Then we show how the 2SD model can account for the eye movement data from two monkeys presented in Stanford et al. (2010). Further technical details for both models are given in the appendix.

## Accelerated race-to-threshold model

In the <u>accelerated race-to-threshold model</u> (Stanford et al., 2010; Shankar et al., 2011), two competing variables, *x_L_* and *x_R_*, denote the developing motor plan to perform a saccade to one of the two targets (in neurophysiological terms, the activity of neurons that trigger eye movements to the left or right), and the variable reaching a certain fixed threshold first (the winner of the race) determines the direction of the saccade that occurs a short efferent delay later. Two different stages must be distinguished: one in which no cue information is yet available, and another, in which the target cue boosts one of the motor plans and suppresses the other. This information is incorporated into the model by assuming build-up rates *r_R_* and *r_L_* of *x_L_* and *x_R_*, respectively, drawn from a (truncated) bivariate Gaussian distribution; if one of the variables reaches a threshold during this stage, the outcome is a coin toss because the buildup rates were sampled randomly from a symmetric distribution. Otherwise, the two oculomotor plans keep changing until the cue information arrives. Once the cue arrives showing, e.g., that the target is on the right-hand side, the build-up rate of *x_R_* increases and and that of *x_L_* decreases (again at constant rates) until a threshold is reached. Fig 2 depicts the trajectories of 4 developing motor plans (hypothetical trials) for the race-to-threshold model: 2 with a gap of 50 ms (**A** and B) and 2 with a gap of 250 ms (**C** and **D**).

**Figure 2.**
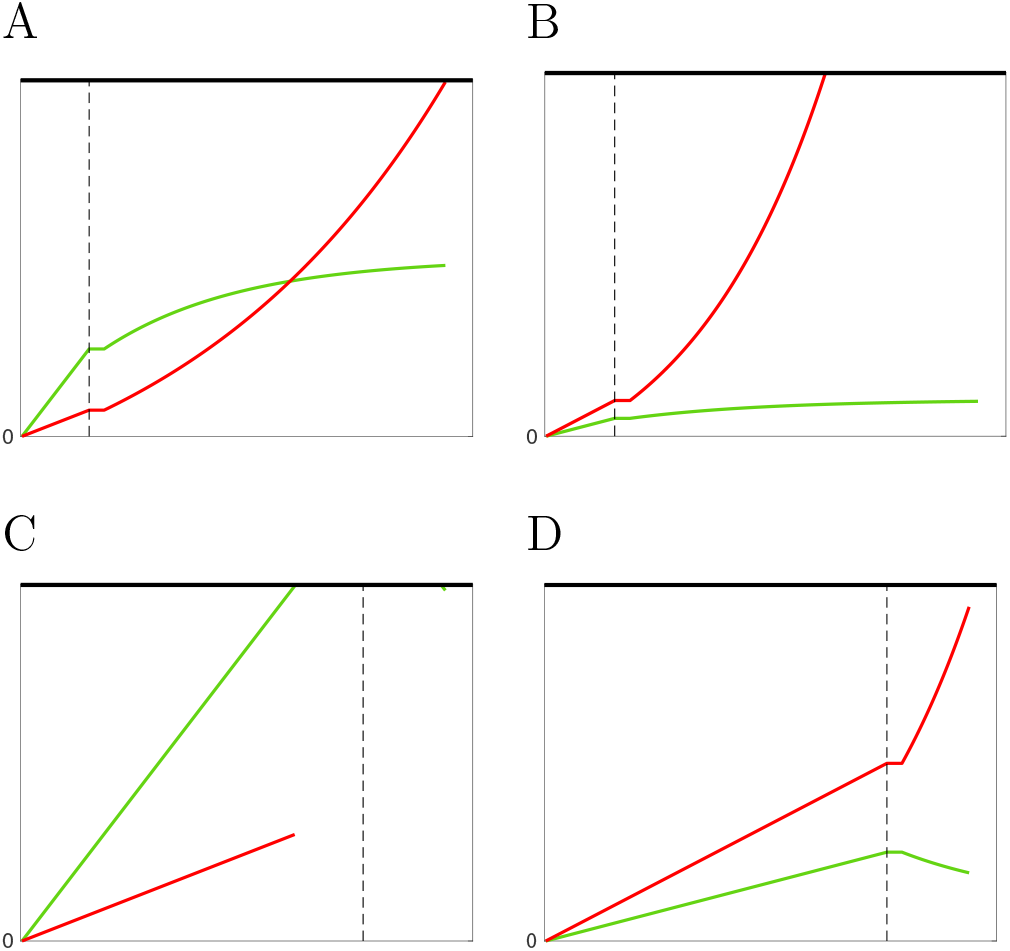
Accelerated race-to-threshold model. Hypothetical trajectories of 4 developing motor plans for the race-to-threshold model: 2 with a gap of 50 ms (**A** and **B**) and 2 with a gap of 250 ms (**C** and **D**). In the first stage, buildup rates are drawn from a (truncated) bivariate Gaussian distribution

## The two-stage diffusion model

The two-stage diffusion model, like the accelerated race-to-threshold model, assumes that the decision is driven by two sub-processes, the first stage feeding directly into the second. The diffusion model is conceived as a stochastic process, *X*(*t*), representing the numerical value of the accumulated “evidence” at time *t* with drift rate, *μ_i_*(*x, t*)(*i* = 1, 2), changing from the first to the second stage. Note that, here, “evidence” is conceived of as an abstract concept and may have different interpretations (see below).

The first stage represents the process from onset of the go signal to the cue signal (the *gap time*). During the gap time, the direction of the trajectory may randomly change from right to left and *vice versa*. Because no target evidence is yet provided, the probability to make an eye movement to the right or left is the same, i.e. *μ*_1_(*x, t*) = 0. With increasing gap time, the probability to reach one of the criteria for making a right or left movement increases (Fig 3). Thus, while the drift rate is zero, the latter fact allows interpreting *X*(*t*) in the first stage as representing an (increasing) urgency-to-respond signal.

**Figure 3.**
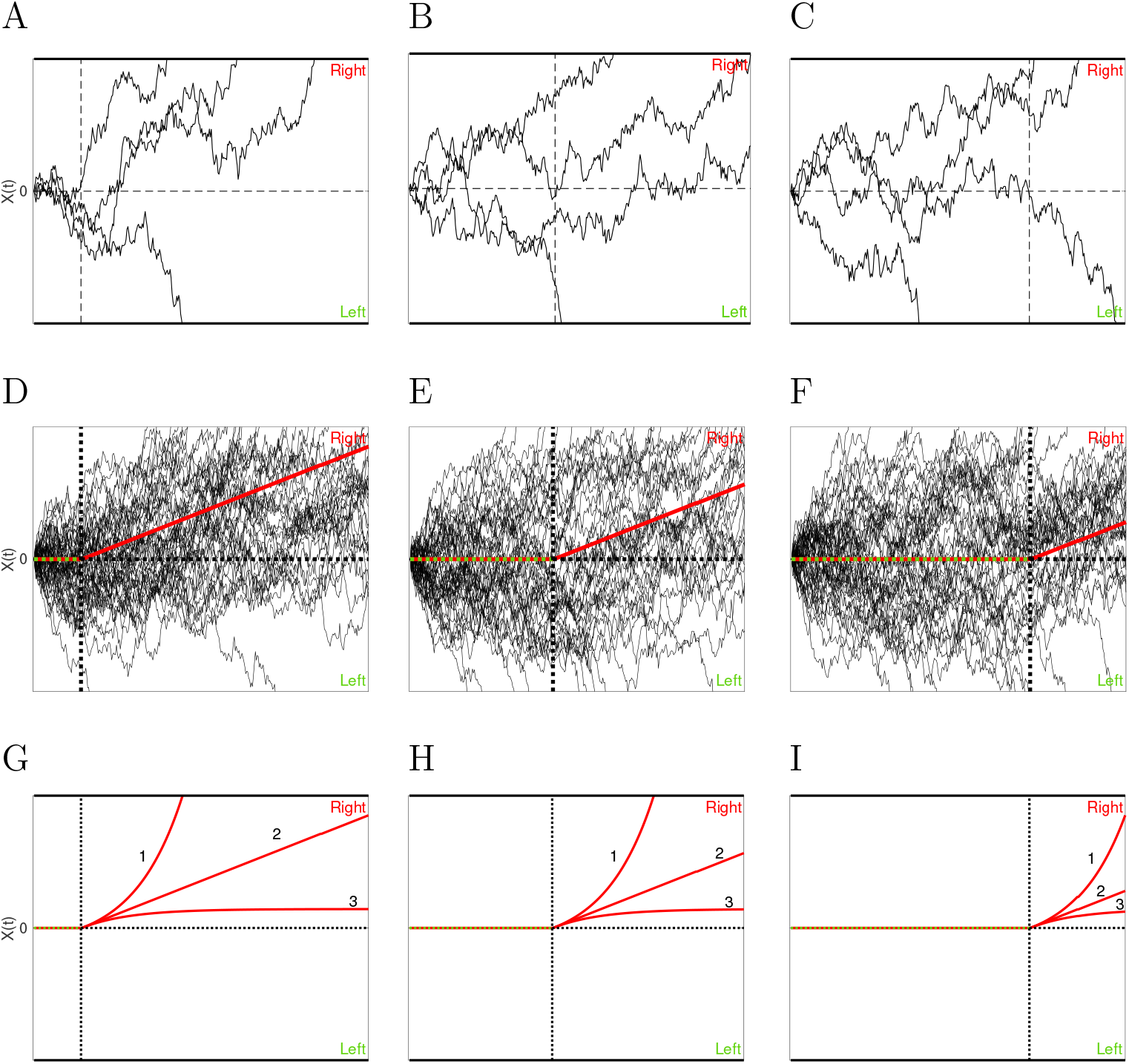
Two-stage-diffusion model. For these examples, the target is red, located on the right-hand side. Each trial is presented as a trajectory from a stochastic process *X*(*t*). **A** – **C** show 4 trials with 3 different gap times [ms]: 50 (**A**), 150 (**B**), and 250 (**C**), indicated by vertical dashed lines. Once a trajectory crosses a threshold (upper and lower horizontal bold face lines) a response is initiated. The upper boundary is associated with the right response, the lower one with a left response. In **A**, 3 out of 4 trajectories cross the upper threshold and, therefore, 3 correct responses are initiated. In **B**, two trajectories cross the thresholds and a correct and an incorrect response are initiated. From the figure it is not clear yet where the 2 remaining trajectories would end up. In **C**, one response occurs during the gap (trajectory hits the lower threshold) and in this example, an incorrect response is initiated. Of the remaining 3 trials, 2 are correct responses (hitting the upper threshold) and 1 is incorrect (hitting the lower threshold). **D** – **F** show the same scenario with 50 trials each. In **D**, showing a short gap time, the vast majority of trajectories are absorbed at the upper threshold and, therefore, correct responses are initiated. In **F**,

The second stage encompasses the process from cue onset to saccade initiation. For a given trial, the accumulation process of the second stage starts at the level where the process of the first stage ended, continuing now with a drift rate *μ*_2_(*x, t*), different from zero. Depending on gap time length, the first process may be closer or farther away from the decision criterion (threshold), and the direction of the trajectory in the second stage may be reversed with more or less success. Once the cumulative evidence exceeds a threshold criterion, the process stops and a response is initiated. Let us consider the behavior of this stochastic process in more detail.

### Different scenarios for the trajectories in the two-stage diffusion model

In this example, the target is a red dot presented on the right, a correct response is initiated as soon as the upper threshold is crossed. Thus, the upper threshold is associated with response *R* (looking to the right), the lower one with response *L* (looking to the left).

Fig 3 shows the basic ideas of the model with hypothetical trajectories. The decision process for each trial is presented as a trajectory generated from the stochastic process *X*(*t*). Assuming that the target is red and presented on the right, a correct response is initiated a soon as the upper threshold, *θ >* 0, is crossed: *X*(*t*) = *θ*. Alternatively, as soon as the accumulated evidence reaches the lower criterion value (here, *X*(*t*) = *θ <* 0), the incorrect response *L* is initiated.

**A** – **C** of Fig 3 show 4 trials with 3 different gap times [ms]. **A**: 50, **B**: 100, and **C**: 250, indicated by the vertical dashed lines. At that time point, the process switches from stage 1 to stage 2. Once a trajectory crosses a threshold (upper or lower horizontal bold face lines) a response is initiated. In **A**, 3 out of 4 trajectories cross the upper threshold and, therefore, 3 correct responses are initiated. Note that even if the evidence is overwhelmingly strong (when the cue is presented rather soon after the go signal), there is a small chance for making an incorrect response. This is an integral part of the stochastic processing mechanism and may be due to attentional lapses. In **B**, one trajectory crosses the upper, another crosses the lower threshold and a correct and incorrect response is initiated, respectively. From the figure, it is not visible where the 2 remaining trajectories would end up, i.e. initiating an *R* or *L* response. In **C**, one response occurs during the gap time (trajectory hits the lower threshold) and, in this case, an incorrect response is initiated. Of the remaining 3 trials, 2 are correct responses (hitting the upper threshold) and 1 is incorrect (hitting the lower threshold).

**D** – **F** of Fig 3 illustrate the same scenario with 50 trials each. The simulated trajectories show that with an increasing number of trials, the overall process converges towards specific directions. In **D**, with a short gap time, the vast majority of trajectories are absorbed at the upper threshold and most of the initiated responses are correct. The reason is that the information about the target occurs almost immediately after the go signal, so an incorrect direction can more easily be reversed. In **F**, with a long gap time, about half of the trajectories are absorbed at the upper or lower threshold. That is, with long gap times, the chances to make a correct or an incorrect response are about the same. A redirection of the movement is almost impossible. The bold lines in **D** – **F** indicate the expected (mean) values of all trajectories. From the onset of the go signal up to the cue, the expected value is the same for making a movement to the right or left side, i.e. the drift rate of the process is 0 (red-green line). After the target color has been revealed, the chances for making a correct response increase and the drift rate, for this example, becomes positive (red line).

**G** – **I** show the mean drifts (expected value) for 3 different model versions after the target appearance. Stage 1 is the same for all versions and described by a Wiener process with drift rate *μ*_1_(*x, t*) = 0. After the target color has been revealed (in the example, red) – stage 2 – evidence accumulation for making an *R* response is directed toward that threshold. First, consider the mean drift case 2 that assumes a Wiener process with drift rate *μ*_2_(*x, t*) = *δ >* 0, which is identical to the one in **D** – **F**: the expected (mean) evidence increases linearly as a function of time.

A widely accepted assumption in diffusion modeling is that the drift rate mirrors the quality of the stimuli (e.g., Ratcliff et al., 2016): the better the quality, the larger is the drift rate (slope). For instance, when the target and distracter colors are difficult to discriminate (e.g. yellow and orange instead of red and green), then the drift rate is assumed to be smaller and the probability for a correct response decreases for all gap times.

Stanford et al. (2010) argue that, after the target color has been revealed, the process for making a correct response accelerates and the one for an incorrect slows down. The 2SD model allows to incorporate acceleration/slowing down in processing in stage 2 depending on the current state by modifying the drift rates to *μ*_2_(*x, t*) = *δ −* (*−γ*)*x* and *μ*_2_(*x, t*) = *δ − γx*, respectively, and shown in Fig 3 **G** – **I**, case 1 (acceleration) and case 3 (slowing down). The processes are instances of the Ornstein-Uhlenbeck process (OUP) (for details see e.g., Diederich, 1992; Ricciardi, 1977).

### Predictions

The two-stage diffusion model allows us to derive several qualitative and quantitative predictions from the above definitions and assumptions.

1. The probability for making a correct response decreases as the gap time increases. As shown in Fig 3 for very short gap times, information about the target is revealed almost immediately after the go signal leading to a high probability of making a correct response. The longer the gap the more trials have either been finished during the gap time, i.e., guessing about the correct side, or the process is too close to one threshold so that reverting incorrect direction is no longer possible, i.e., also leading to chance level. Fig 4 shows the predictions of a Wiener process in stage 2 for a large range of parameter values *μ*_2_, gaps, and three different threshold i.e. **A**: low, **B**: medium, and **C**: high. The same pattern occurs when assuming an OUP in stage 2 (not shown).
2. The mean choice response time for incorrect responses is shorter than for correct responses (fast errors). During the gap time no information about the target is yet available. Correct and incorrect responses during this time are random with probability 0.5. After the target is presented information for a correct response is available. That is, when an incorrect response occurs it needs to be fast before new evidence for a correct response is provided. When responses are made only during gap time, i.e. for long ones, the mean choice response times for correct and incorrect responses are identical. Fig 4 (**D** – **I**) shows the predictions assuming a Wiener process in stage 2 for a large range of parameter values *μ*_2_, gaps, and three different thresholds. (**D** – **F**) refer to mean choice times for correct response; (**G** – **I**) for incorrect responses. The same pattern occurs when assuming an OUP in stage 2 (not shown).
3. The probability distributions are right skewed and bimodal. Furthermore, the overall pattern of the probability distributions is the same regardless of the specific process (Wiener, OUP) of stage 2. Fig 5 shows predicted distributions for the three versions of the model introduced above as a function of gap time. To make them comparable (and in accordance with Stanford et al. (2010)) they are unconditioned (sum of both areas adds up to 1) and normalized (all values are divided by the largest value). Assuming a slowing down depending on the current state in stage 2 obviously predicts slower RT (left column), while an acceleration in stage 2 predicts faster RTs (right column) as compared to a linear increase in evidence (middle column). When all responses are made within the gap time (last row), the process is driven only by stage 1, for which we assumed a standard Wiener process. Thus, the distributions for correct and incorrect response are identical, thus mean RTs are also identical and the probability to choose R or L is .5.

**Figure 4.**
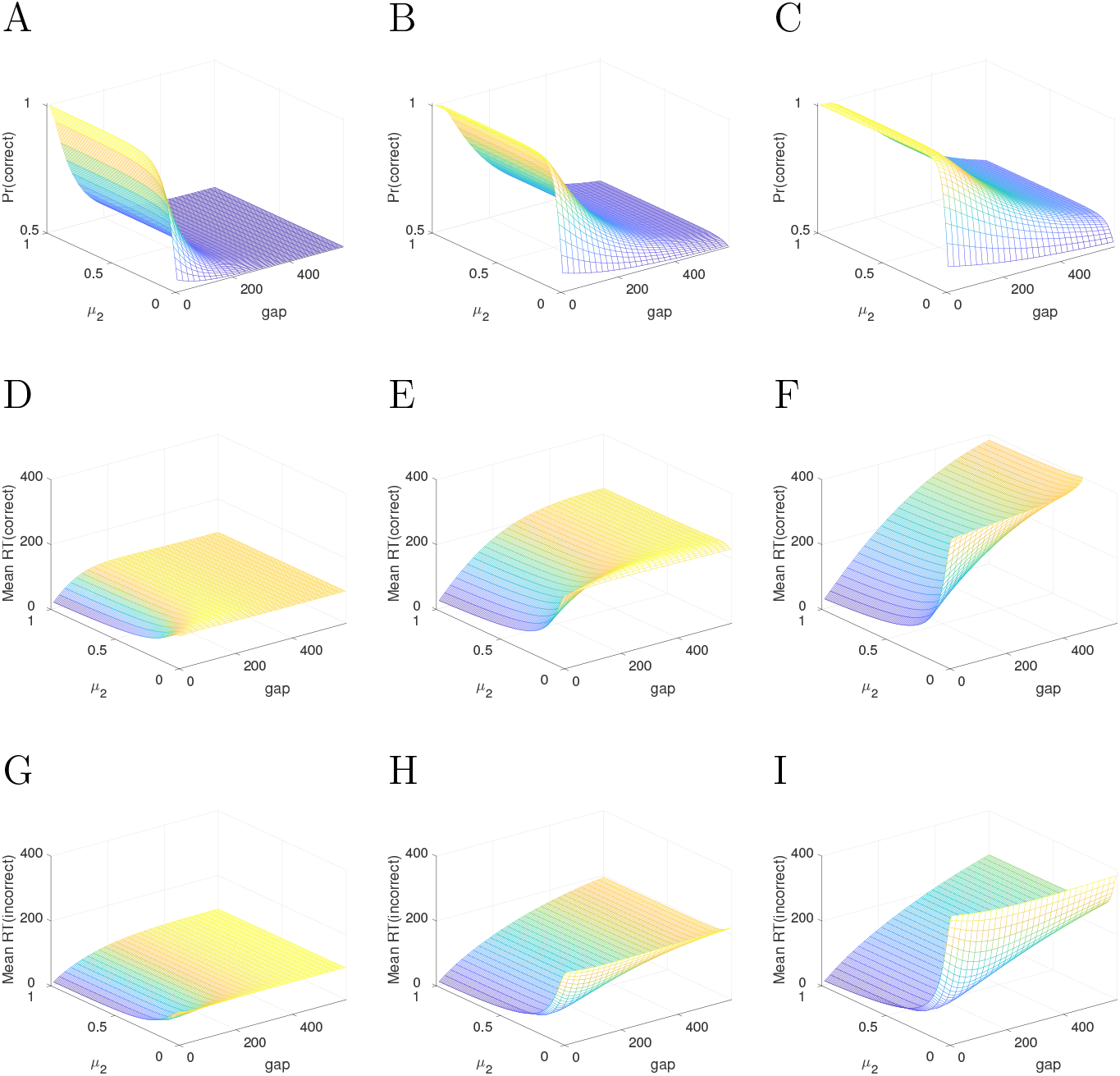
Predicted choice probabilities and mean choice RT. **A** – **C**: Predicted choice probabilities for a correct response as a function of *μ*_2_ and gap times and three different thresholds. **A**: low; **B**: medium; **C**: high. **D** – **F**: Predicted mean choice response times for correct responses as a function of *μ*_2_ and gap times for the respective three different thresholds. **G** – **I**: Predicted mean choice response times for incorrect responses as a function of *μ*_2_ and gap times for the respective three different thresholds.

**Figure 5.**
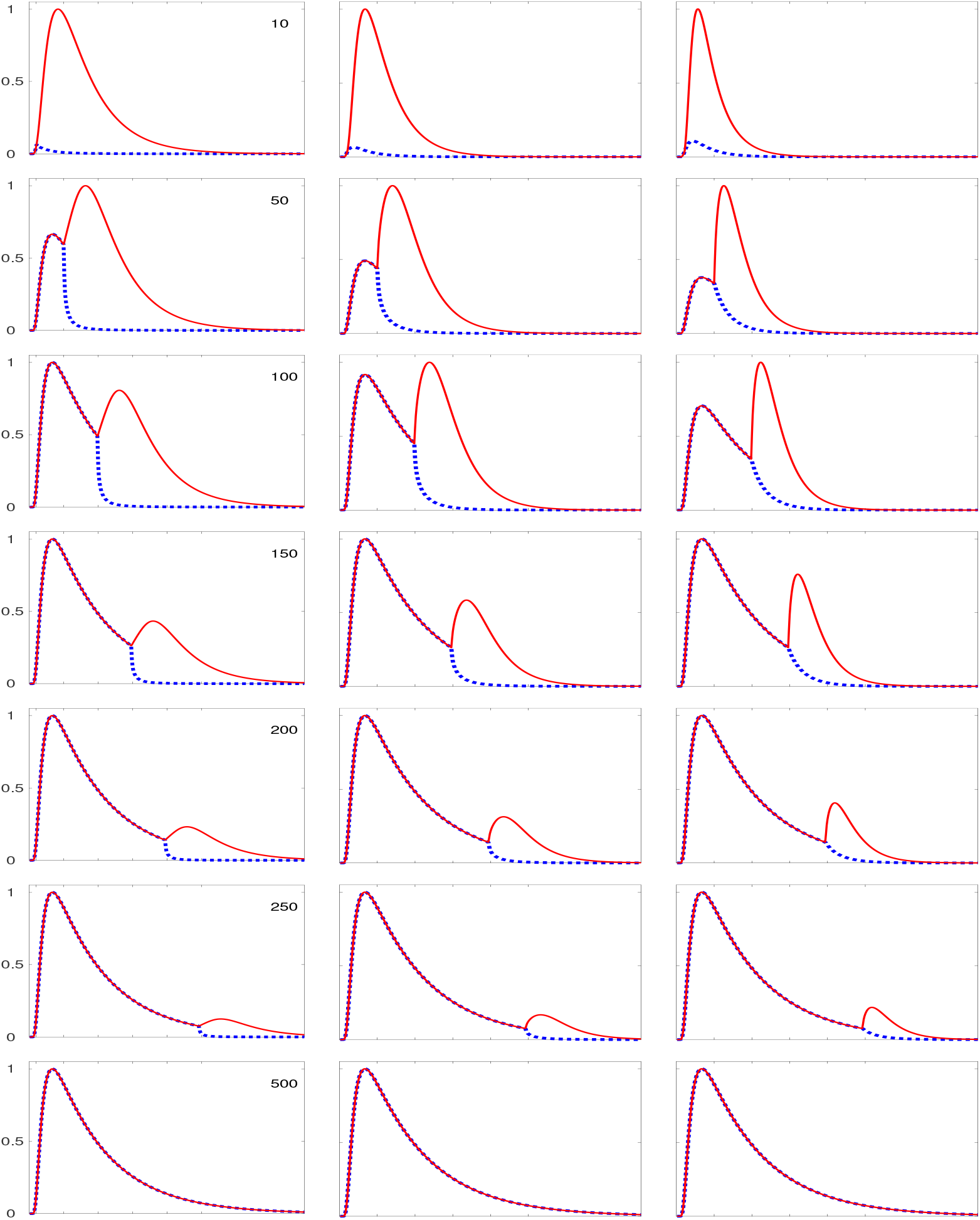
Predicted probability distributions. Each column shows the predicted RT probability distribution (correct: red, incorrect: blue) for one model version for different gap times. Left: OUP with *μ*_2_(*x, t*) = .2 *− .*02; center: Wiener with *μ*_2_(*x, t*) = .2; right: OUP with *μ*_2_(*x, t*) = .2 + .02. As in Shankar et al. (2011), distributions are unconditioned (with respect to correct/incorrect) and normalized (division by maximum value).

### Some extensions

The basic model with two or three parameters (Wiener: drift rate *δ*_2_ for stage 2 and threshold *θ*; *δ, γ* and *θ* for OUP) can be extended to account for experimentally induced effects and other theoretically motivated variations.

1. Biases towards the left or right may be reflected in the starting point of the process or in the drift rate of stage 1. Instructions or rewards may influence the accumulation process. A tendency to look to the right, for instance, may be incorporated by shifting the starting position *X*(0) closer to the right-hand threshold (in the current example, *X*(0) *>* 0). Similarly, a tendency to look to the left is reflected by assuming *X*(0) *<*. In this case, the bias may wear off when the target appears late (long gaps). Assuming that the bias is incorporated into the drift rate, i.e. *μ*_1_(*x, t*) ≠ 0 the bias becomes stronger with longer gaps (Diederich, 2008; Diederich & Busemeyer, 2006).
2. The gap time may not be fixed but may follow a distribution. For the basic model, we assume that the switch from stage 1 to stage 2 occurs exactly at the respective gap time point. However, it is possible that the switch is delayed and/or follows a probability distribution. Diederich and Oswald (2014) investigated several distributions for the switching time, among them geometric, Poisson, uniform. The exact shape of the distributions changes whereas, however, the overall pattern does not.
3. The threshold may not be constant but change as a function of gap time. The basic model assumes a constant threshold. However, it is possible that with increasing gap time the threshold cave in (or out). For the two-stage diffusion model, Diederich and Oswald (2016) derived prediction assuming various functional forms for the thresholds and, most importantly and different from all other models, with incoming boundaries, separately for stage 1 and 2. Also note that the threshold may be related to the available time to make a decision. That is, the threshold is assumed to be a function of deadlines with smaller thresholds for time constraints (Diederich & Oswald, 2016).

At the moment, we do not consider these model extensions any further, but they could be incorporated if physiological evidence require it and if corresponding data to test those assumptions were available.

## Two-stage diffusion model account of data from Stanford et al. 2010

In the following, we show how the basic two-stage diffusion model accounts for the data from Stanford et al. (2010) (data for monkeys S and G, provided by courtesy of Emilio Salinas).

Typically, the response time, *RT*, is assumed to be the sum of two random variables, i.e., the decision time *T_D_*, here modeled by the diffusion model, and a non-decision time *T_ND_* (e.g., Luce, 1986):

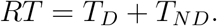

We do not make any assumptions about the distribution of *T_ND_* here, simply setting it to a constant. Thus, the basic two-stage diffusion model has only three parameters to be estimated from the data: drift rate *μ*_2_, threshold *θ*, and *T_ND_* (note that *μ*_1_ = 0 *a-priori*).

The model is implemented as a continuous Markov process approximated by a discrete Markov chain (for the matrix approach, see Diederich, 1997; Diederich & Busemeyer, 2003; Diederich & Oswald, 2014, 2016). The drift coefficient was set to *σ* = 1; the time unit to *τ* = .1 (that is, 10 sample points per 1 ms). The resulting step size of the process is 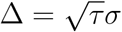. The step size reflects the amount of evidence sampled per time unit (for details, see appendix).

Importantly, in testing the model we first estimate the parameters from a subset of the (smoothed) data for some of the gap conditions and then predict the remaining gap conditions as well as mean choice response times and choice probabilities. In particular, we estimate the parameters using maximum likelihood estimation and utilize the Matlab routine *fminsearchbnd*, which allows parameters restrictions. It is similar to *fminsearch* that uses the Nelder-Mead simplex search method.

Fig 6 and Fig 7 show the data and model accounts for monkeys S and G, respectively. To produce the empirical distributions in **A** and **C** of Figs 6 and 7, the observed response times were arranged in bins of 4 ms and then smoothed with a moving average filter with filter length *M* = 20 (see Stanford et al., 2010). Parameters were estimated from gap conditions 50, 100, 150, 200, and 250 ms (**A**) and the respective model account is shown in **B**. With these parameter estimates, the model predicts the patterns **D** for gap conditions 75, 125, 175, and 225 ms for data **C**. Furthermore, with the same parameters the model predicts choice probabilities **E** and mean choice response times **F** for correct and incorrect choices. The model gives a good qualitative account of the data. Clearly, the data show bimodal distributions as a function of gap times and faster mean choice response times for incorrect responses (fast errors) as predicted by the model. Quantitatively, the model overestimates/underestimates for some gap conditions. Given that only three parameters were estimated from about half of the data, the model is obviously able to account for the data.

**Figure 6.**
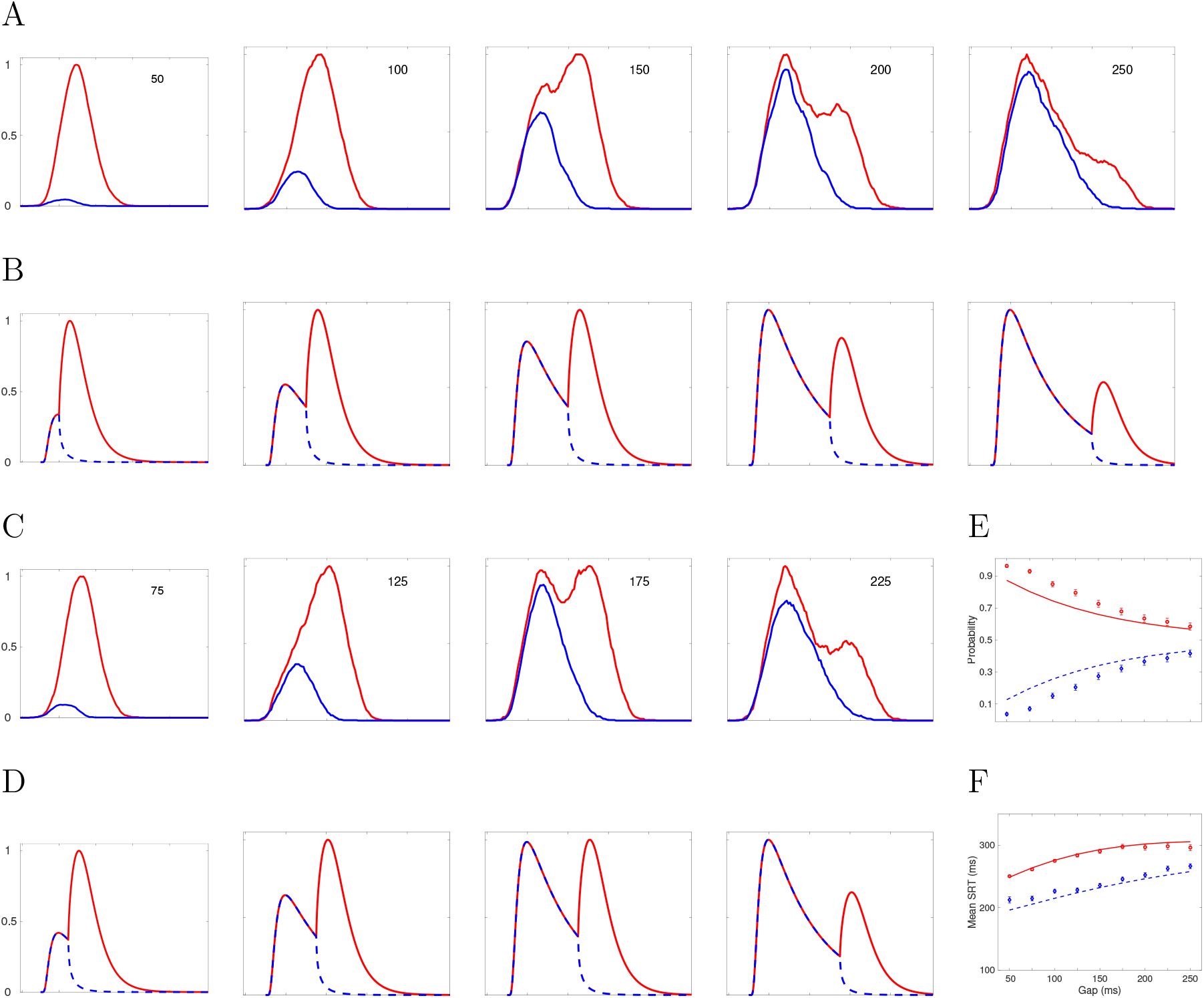
Data and model account for Monkey S. **A** shows the smoothed empirical distributions for correct (red) and incorrect (blue) response times for gap conditions 50, 100, 150, 200, and 250 ms and **C** those for gap conditions 75, 125, 175, and 225 ms. **B** and **D** depict the corresponding predictions from the two-stage diffusion model. **E** and **F** show probabilities and mean RT data (points including 95% confidence intervals) and model predictions (lines) (red correct, blue: incorrect responses). All parameters were estimated using data from the upper row of **A** only with values *μ*_2_ = 0.3, *θ* = 13, and *T_ND_* = 160 ms.

**Figure 7.**
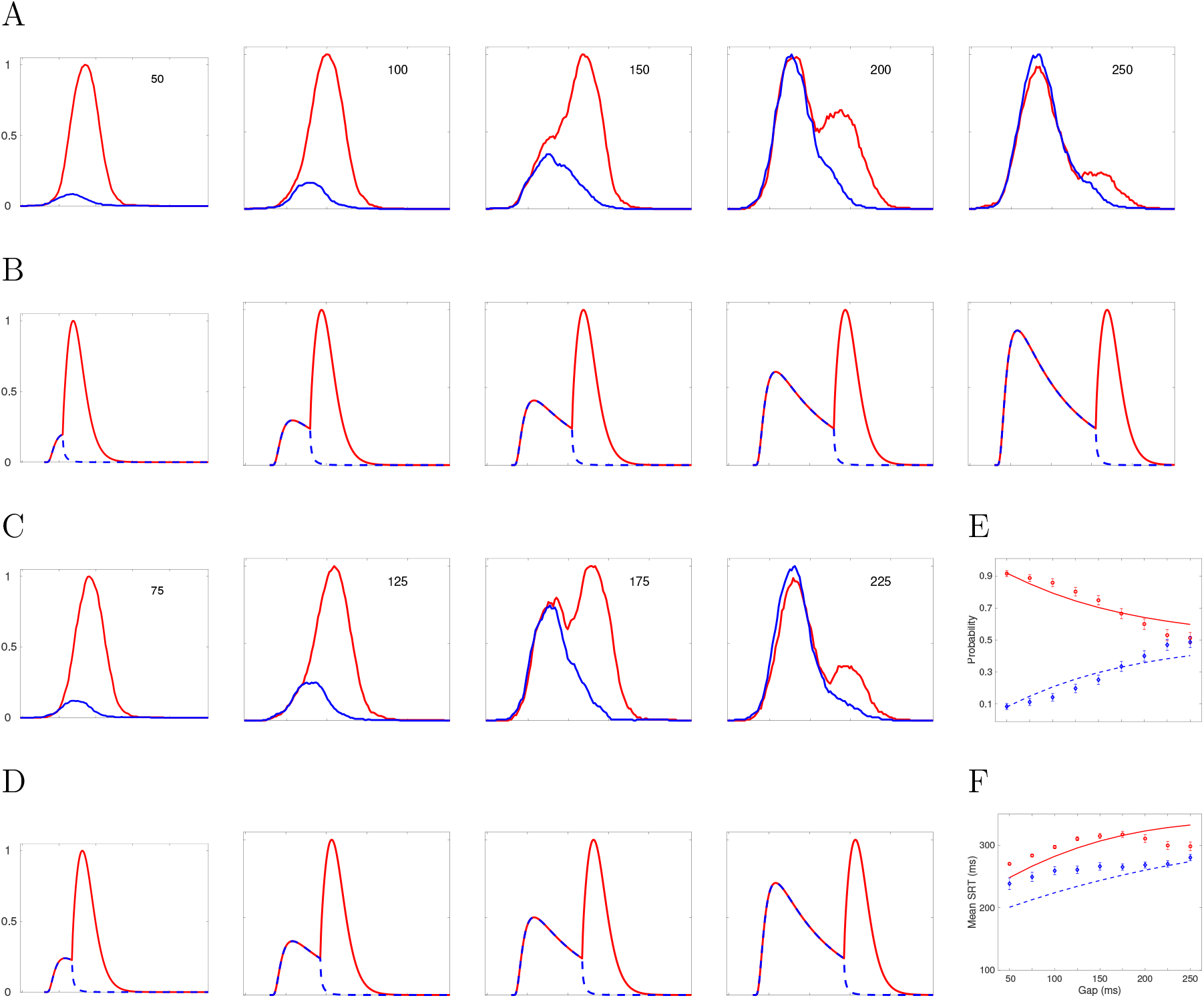
Data and model account for Monkey G. For the description see Fig 6 for Monkey S. Parameters for monkey G are *μ*_2_ = 0.2, *θ* = 12, and *T_ND_* = 150 ms.

## Discussion

The compelled-saccade task, introduced in Stanford et al. (2010) and further analyzed in a number of eye movement studies (e.g., Costello et al., 2013; Salinas & Stanford, 2013; Salinas et al., 2014; Salinas & Stanford, 2018; Shankar et al., 2011), has been very successful in tracking perceptual performance. Salinas and colleagues have developed a simple, heuristic model that is able to reproduce most of the observed behavior while separating perceptual from motor processing components. As observed by Drugowitsch and Pouget (2010), however, this accelerated-race-to-threshold model does not square with the most successful theoretical approach for decision making under time pressure, i.e. stochastic diffusion models (e.g., Ratcliff et al., 2016). Specifically, in the former the primary source of random variability is in the initial state of information buildup across trials, whereas in diffusion-type models variable sensory information is accumulated randomly fluctuating over the entire duration of the trial. Thus, the accelerated-race-to-threshold model implies that incorrect decisions are due to the inertia of the racing variables that have started with an across-trials randomly sampled constant buildup in the “wrong” direction^1^.

Here, we have shown that a two-stage-diffusion model is able to account for the rather complex pattern of data generated in the compelled response task: with only 3 easy-to-interpret parameters, the model is able to predict both response time distributions and choice probabilities for arbitrary gap time conditions in the compelled-response task. It is important to realize, however, that an important feature of the two-stage model proposed here is that it is a non-time-homogeneous stochastic process, in contrast to the “common” drift-diffusion model (e.g., Ratcliff & Tuerlinckx, 2002). This non-homogeneity property is essential in representing the effect of the cue on the on-going saccade trajectory. Although the model is mathematically complex, it is analytically tractable using the discrete Markov chain approach developed by Diederich and colleagues (Diederich & Oswald, 2016; Diederich, 1992, 1997; Diederich & Busemeyer, 2006) that permits us to derive predictions without having to rely on model simulations.

Another feature that distinguishes the two-stage model from both the race-to-threshold and the drift-diffusion model is that there is no need to assume between-trial variability to match the pattern of data across conditions, i.e. all parameters remain constant from one trial to the next rather than being sampled from some additionally-assumed probability distribution. This is an important property in light of the recent criticism by Jones and Dzhafarov (2014b) arguing that across-trial variability make models overly flexible(see also Heathcote, Wagenmakers, & Brown, 2014; Smith, Ratcliff, & McKoon, 2014; Jones & Dzhafarov, 2014a). The two-stage diffusion model is more than a data-fitting tool: it is eminently testable and is falsifiable already at the level of qualitative features, for example in predicting the bimodality of the RT probability distributions for particular gap times and, most importantly, fast errors.

The race-to-threshold model produces a good fit to various data sets from the compelled-response paradigm and quantitatively accounts for the separate components of the task: motor response, perceptual evaluation, and additional cognitive factors. However, it does so by introducing a number of auxiliary assumptions. For example, Shankar et al. (2011) assume that the gap time is normally distributed with a mean of the experimentally administered gap time and a variance identical to the one of the non-decision time. For short gap times, this leads to negative gap times; but eliminating negative gap times would result in a distribution that is no longer Gaussian. In addition Shankar et al. (2011) assume that the transition from stage 1 to stage 2 is interrupted for some time. That is, the process is constant for some time before acceleration or deceleration occurs. How long this interruption lasts is determined by two parameters to be estimated from the data. The probability of “confusion” is another parameter introduced ad-hoc to account for lapses, i.e., trajectories going into the wrong direction even for short gap times due to inattention or something else. It is required in the race-to-threshold model because the process is deterministic during one trial. Note that, on the other hand, in a stochastic diffusion process model this is built-in into the process itself. Finally, Stanford et al. (2010) and Shankar et al. (2011) assume a parameter called sensory discrimination time, that is the time to reach the final values of the direction changers for target and distracter. How this parameter is to be interpreted – given its large range (Monkey S 85 ms; Monkey G 1800 ms) – is not intuitive. Altogether, the race-to-threshold model requires 11 parameters to achieve a good fit to the data. While these additional parameters lead to an improved quantitative fit, we feel that the model is conceived of as a measurement device rather than a model explaining the underlying decision process.

Note that the two-stage diffusion model’s fit could be improved upon by including some of the assumptions stated in the appendix and by specifying the non-decision time further. Instead of assuming a constant non-decision time, *T_ND_* may be considered a random variable. Likewise, Stanford et al. (2010) assume a normal distribution and Ratcliff et al. (2016) a uniform distribution, but the choice of a particular distribution is mainly guided by mathematical tractability rather than by behavioral or physiological evidence. Behavioral data will not be able to solve this problem, however (for an early discussion see Luce, 1986). To sum up, we propose the two-stage diffusion model as an alternative to the race-to-threshold model for the compelled-response task. This note demonstrates that the data pattern from this task can be described and predicted by a model that accumulates randomly fluctuating sensory information within each trial, without the need for simulation and with only a few parameters to be estimated. While we do not take up a stance on which model type is better supported by neurophysiological data, there is ample evidence that stochastic-diffusion type models provide a quantitative link between the time-course of behavioral decisions and the growth of stimulus information in neural firing data (see, e.g., Smith & Ratcliff, 2004).

## Acknowledgments

We are most grateful to Emilio Salinas, Terry Stanford, and colleagues for very helpfully providing us with their data and programs.

## Appendix Further model details

### Race-to-threshold model

According to Shankar et al. (2011), in a given trial both variables *x_L_* and *x_R_* start with a value of 0 and race to a threshold of 1000 units. Build-up rates *r_R_* and *r_L_* are sampled from a truncated bivariate Gaussian distribution with mean *r_G_* (same for both variables), standard deviation *s_G_* (same for both variables) and correlation coefficient *r*. The first part of the race encompasses the time from the go screen (offset of the fixation point) until the cue screen; the second part from the cue screen until a response it made. Because no information is given in part 1, the winner is random with probability 0.5. In part 2, after the color has been revealed, the racer with the “correct” color direction – the target – accelerates and the one with the “incorrect” color direction – the distracter – slows down.

**Stage 1:** For a given trial, the rates (change as a function of time)

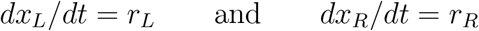

are constant and linear but vary across trials.

**Stage 2:** Let 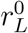 and 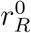 be the buildup rates drawn initially; after the target color is revealed, the rates in stage 2 change. The competing variables start accelerating according to the locations of the target and distracter. If the target is on the right side, then the buildup rate of *x_R_* approaches a large, positive value *r_T_* (for target) and the buildup rate of *x_L_* approaches a small or negative value *r_D_* (for distracter). The corresponding equations are

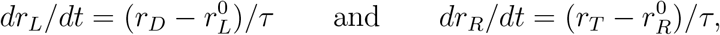

with the added rule that once the buildup rates reach their new target values–that is, once *r_L_* is equal to *r_D_* and *r_R_* is equal to *r_T_* – they stop changing, so the last two derivatives become zero. In this way, the acceleration is constant but lasts a finite amount of time, which is precisely equal to *τ*, and *r_T_* is the maximum possible buildup rate for the variable that generates a movement toward the target (see Shankar et al., 2011).

When the target is to the left of fixation, the roles of *x_L_* and *x_R_* are reversed: when the sensory information arrives, *r_L_* increases toward the high rate *r_T_* and *r_R_* decreases toward the low rate *r_D_*, so

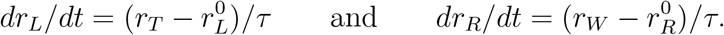

Several more parameters are included:

- *p_e_* is the probability of confusion, accounting for lapses. That is, the monkey has enough time to make a correct answer but failed to do so.
- The non-decision time (collapsing afferent and efferent processing times) is drawn from a Gaussian distribution withe mean *T_ND_* and standard deviation *σ_ND_*.
- The start and end of an interruption time *t* = *I*_1_ and *t* = *I*_2_ from stage 1 to 2 during which no change in the builup rates occurs.

Altogether, the model includes 11 parameters to be estimated from data using simulations and extended grid search. In addition, some further fixed parameters are included in the simulation procedure. Moreover, in the program the gap time is not fixed but follows a normal distribution, *N* (*gap, σ_ND_*).

### Two-stage diffusion model

The decision process is modeled as a two-stage diffusion model, with stochastic process *X*(*t*) representing the numerical value of the accumulated evidence at time *t*. Note that, as explained above, “evidence” is to be interpreted as an abstract notion depending on the context/stage of accumulation: upon the offset of the fixation point, the decision maker sequentially samples information from the stimulus display over time, retrieves information from memory, or forms preferences, depending on the context.

The small increments of evidence sampled at any moment in time are such that they either favor option left (*L*) (*dX*(*t*) *>* 0) or option right (*R*) (*dX*(*t*) *<* 0). The evidence is incremented according to a diffusion process:

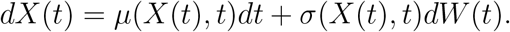

Here, *μ*(*x, t*) is called the *effective drift rate* describing the instantaneous rate of expected increment change at time *t* and state *x* = *X*(*t*). The factor *σ*(*x, t*) in front of the instantaneous increments *dW* (*t*) of a standard Wiener process *W* (*t*) is called the *diffusion coefficient*, and relates to the variance of the increments.

Here we assume that the process in the first stage is a Wiener process *W* (*t*) with drift rate *μ*(*x, t*) = *μ*_1_ = 0 and diffusion coefficient *σ*(*x, t*) = *σ*_1_, i.e.

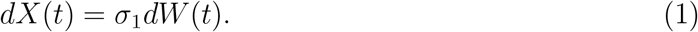

Because no evidence is given in the first part (from fixation offset and Go screen to the target), the drift rate in the first stage is assumed to be 0, so that it is equally likely to make saccade to the left or to the right side.

With the appearance of the cue screen, target information is revealed, and the second process starts. Depending on the color of the fixation point, a movement to the right side is correct or incorrect (similar for the left side). This part is modeled as a Wiener process with drift *μ*(*x, t*) = *μ*_2_ and diffusion coefficient *σ*(*x, t*) = *σ*_2_, i.e.

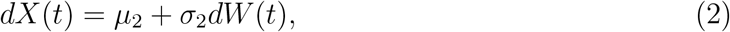

or, with *μ*(*x, t*) = *δ −γ x* and *σ*(*x, t*) = *σ*_2_, i.e.

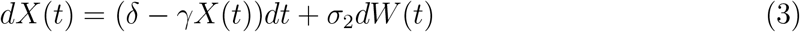

as an Ornstein-Uhlenbeck process (OUP). Setting *γ >* 0 yields evidence accumulation towards one of the choice options at a linearly decaying rate, that is, it induces a change of the effective drift rate *μ*(*x, t*) = *δ − γx* depending on the current state *x*. Setting *γ <* 0 accelerates the process. Note, however, that the process might become unstable and, strictly speaking, it is not called OUP for *γ <* 0. Finally, setting *γ* = 0 reduces Eq. 3 to a Wiener process with drift (Eq. 2). Furthermore note that, the entire processes is non-time-homogeneous.

The process continues until the magnitude of the cumulative evidence exceeds a threshold criterion, *θ*. Then, the process stops and response *L* is initiated. Or it stops and an *R* response is initiated if the accumulated evidence reaches a criterion value for choosing response *R* (here, *X*(*t*) = *θ <* 0).

The Wiener process with drift and two absorbing boundaries has four basic parameters (drift rate, diffusion coefficient, rating point, boundary) of which one can be set arbitrarily. Here we fix the diffusion coefficient *σ* which is basically a scaling factor by setting *σ*_1_ = *σ*_2_ = 1 (e.g., Diederich & Mallahi-Karai, 2018). Furthermore, we assume that the participant has no a-priori bias was towards one direction (left or right) and, therefore, we set the starting point of the process to *X*(0) = 0. In this context, if *X*(0) *>* 0, the participant would a-priori favor looking to the left (*L*); for *X*(0) *<* 0, the participant has an a-priori tendency (bias) for looking more often to the right than to the left. This initial state could also be governed by a probability distribution.

Similar to the race-to threshold model, we assume a residual time *T_ND_* and also distinguish between a correct *R* and a correct *L* response. For our purposes, we simply assume *T_ND_* to be a constant because we do not know anything about the underlying distribution. With that, the two-stage diffusion model has only three (Wiener process in stage 2) or four (OUP in stage 2) free parameters in total.

## Footnotes

1 Similar notions of a linear rise-to-threshold have been proposed earlier for typical eye movement tasks with highly detectable stimuli, e.g., by Carpenter and colleagues (Carpenter & Williams, 1995; Carpenter, Reddi, & Anderson, 2009). Moreover, the linear-ballistic-accumulator model, introduced in Brown and Heathcote (2008) as a simplified alternative to diffusion models that also proposes a constant buildup of random slope, has gained some popularity for psychological choice RT paradigms.

